# In situ polymerized monolith tips for reproducible, format-flexible proteomic sample preparation applied to biofluids

**DOI:** 10.64898/2026.06.11.731522

**Authors:** Kathrin Korff, Lukas Henneberg, Denys Oliinyk, Nils Eikmeier, Anton M. Schüle, Tim Heymann, Alicia-Sophie Schebesta, Vincent Albrecht, Christoph J.O. Kaiser, Matthias Mann, Johannes B. Müller-Reif

## Abstract

Sample preparation increasingly sets the throughput and reproducibility of mass spectrometry (MS)-based proteomics. StageTips (stop-and-go extraction tips) and variants thereof have long been common implements to purify samples, and we recently extended the concept to solid-phase extraction capture (SPEC) tips, in which the entire digestion takes place in sub-microliter volumes. Here we replace the hand-packed bed with a strong anion-exchange (SAX) monolith photopolymerized directly inside the pipette tip from a defined recipe (pSPEC). A liquid-handling robot casts 384 tunable tips in minutes, at low cost and in any format, adding negligibly to the workflow’s variance. Across biofluids, pSPEC added ∼20% more identifications than in-solution plasma and reached 3,500 protein groups from a single injection of healthy urine and 4,800 from saliva at 100 samples per day, depths usually requiring depletion or fractionation. The same light-cast chemistry should extend to single cells, affinity capture, and population-scale studies.

## Introduction

Mass spectrometry-based proteomics has matured into a technology for deep, quantitative protein analysis across scales from single cells to clinical cohorts (*1, 2*). Unfolding this potential at scale, however, still requires a clean-up step before injection for liquid chromatography-mass spectrometry (LC-MS) analysis (*3*–*8*). In 2003, our group introduced the StageTip, a pipette tip holding a small plug of chromatographic material that desalts and concentrates peptides at the bench (*3*). Its descendants extended the idea from clean-up to complete sample preparation: in-StageTip (iST) processing performs reduction, alkylation and digestion within a single tip (*8*), filter-aided sample preparation (FASP) performs protein chemistry using the tip as a barrier (*7*), and integrated spin-tip workflows fold capture, reaction and purification into the same confined volume (*9*). The appeal has been that by miniaturizing and confining the chemistry, losses drop, contaminants wash away, and many samples can be handled in parallel (*10*). The Evosep HPLC system is likewise built in part on the StageTip principle (*11*).

We recently pushed this confinement principle further by developing solid-phase extraction capture (SPEC), in which a strong anion-exchange (SAX) stationary phase binds intact protein, sequesters the detergents and salts required for efficient lysis. This supports in-tip digestion within a nanoliter substrate volume, so that the chemistry of the tip itself, rather than a separate depletion or enrichment step, defines what reaches the mass spectrometer (*12*). SPEC yields deep, low-input proteomes from cells, tissues, plasma and other demanding matrices.

Despite its attractions, the upfront capture-and-digest step in SPEC is still laborious, because its performance-defining component, the stationary phase, is built by hand. Packed-bead and disc-based tips require a frit and the resin bed is typically assembled manually; packing is slow, scales only to a few hundred tips, and is operator-dependent, so small differences in bed density alter binding capacity and back-pressure and propagate into the final quantitative result. The reproducibility of this step is thus set not by its chemistry but by the manual operation that builds it.

We reasoned that in situ polymerization might offer a way out: a porous monolith cast and cured directly inside the tip, its geometry defined by the polymerization volume and its chemistry, porosity and capacity set by composition rather than by hand. Monolithic spin-tips prepared by photo-initiated polymerization have been coupled to fast chromatography for proteomics (*13, 14*), establishing the feasibility of such an approach. Here we establish a general principle for proteomic sample preparation, that the performance-defining stationary phase can be a programmable material, specified by recipe and cast in place by light rather than assembled by hand, and demonstrate it as an automated, strong-anion-exchange monolithic SPEC platform. Because the bed is now polymerized in situ from a single recipe optimized jointly for fabrication and proteomic performance, the same chemistry can be produced in 384 tips at a time on a liquid-handling robot within minutes. We show that polymerized SPEC is deeper, faithful and highly reproducible in plasma, and that one chemistry and one protocol generalize across biofluids, reaching state-of-the-art depths in dilute, under-served samples, such as urine and saliva that exceed far more laborious workflows.

## Results

### An automated, in situ polymerized SAX monolith tip platform for SPEC proteomics

The pSPEC platform developed here is built around a strong anion-exchange (SAX) monolith polymerized directly inside a standard pipette tip and used within the solid-phase extraction capture (SPEC) workflow (**Figure 1A**). In SPEC, protein binding, washing, on-tip digestion, and elution all proceed within the nanoliter pore volume of the tip’s stationary phase (Figure 1B). In the two-tip configuration (*12*), lysate is loaded onto the SAX tip, washed, digested, and eluted onto a C18 Evotip for LC–MS, so that the chemistry of the tip itself defines what reaches the mass spectrometer.

**Figure 1.**
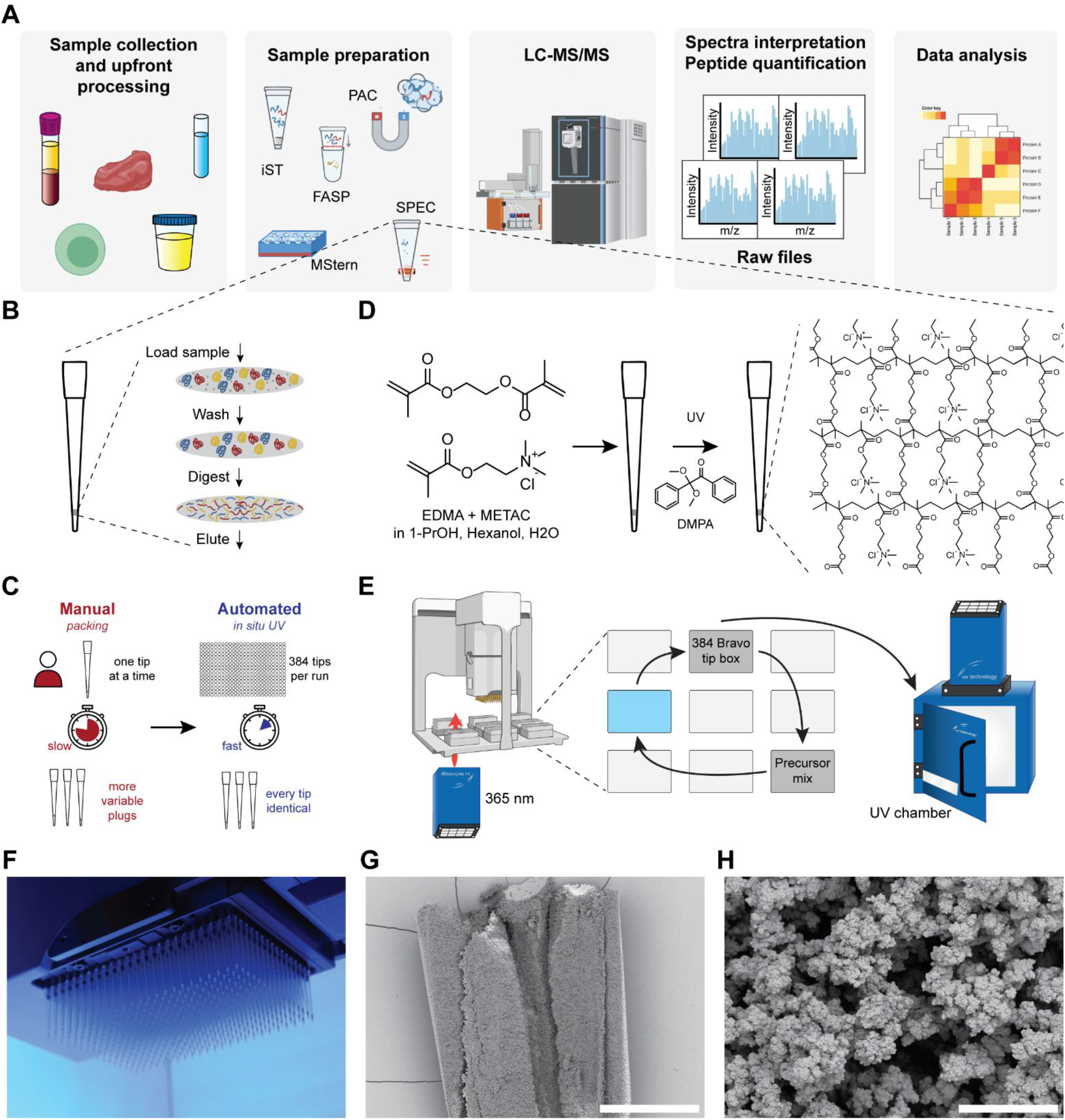
Automated in situ polymerization of SAX monolith tips for solid-phase extraction capture (SPEC) proteomics. (A) SPEC in the context of an LC–MS proteomics pipeline: sample collection and upfront processing → sample preparation (iST, PAC, FASP, MStern, SPEC) → LC-MS/MS → spectrum interpretation and peptide quantification → data analysis. (B) SPEC on-tip cycle: load → wash → on-tip digest → elute onto a C18 Evotip. (C) Manual bead packing versus automated in situ UV polymerization. (D) Monolith chemistry: EDMA (crosslinker) and METAC (SAX functional monomer) with the photoinitiator DMPA in a 1-propanol/hexanol/water porogen, UV-cured into a crosslinked strong-anion-exchange (SAX) network. (E) Robotic fabrication: precursor is aspirated into a 384-tip box and polymerized over a 365 nm UV source, with a final cure in a UV chamber. (F) Photograph of 384 tips during UV polymerization. (G) SEM of a cleaved tip showing the monolith filling the full tip lumen (Aquilos 2, 3.00 kV, ∼50 pA, T1 detector; 191×). Scale bar, 500 µm. (H) High-magnification SEM of the globular, macroporous monolith morphology (Aquilos 2, 20,000×). Scale bar, 5 µm.

As mentioned above, the hand-packed stationary phase has been the workflow’s constraint on scale and reproducibility (*3, 8, 9*) (**Figure 1C**). For pSPEC we form the bed in situ so that the bed’s geometry is fixed by the polymerization volume and its chemistry, porosity, and pore size are set by composition. This allows the stationary-phase composition to be tuned by recipe for each task which we characterize below.

The monolith that we employ is a SAX copolymer of the crosslinker ethylene glycol dimethacrylate (EDMA) and the quaternary-ammonium monomer METAC, whose permanent positive charge supplies the anion-exchange sites at the pH range of interest for proteomics sample preparation (*15*). It is cast from a 1-propanol/hexanol/water porogen with the photoinitiator DMPA and cured under UV (*16, 17*) (**Figure 1C)**. The porogen does not participate in the polymerization but templates the void network and is washed out afterwards, so that monomer ratio, crosslinker content, and porogen composition together set mechanical rigidity, charge density, and pore architecture. Fabrication is automated on a robotic system (here an Agilent Bravo) with a 384-channel head and uses unmodified polypropylene Bravo tips (**Figure 1D**): the head aspirates ∼0.5 µL of precursor into all 384 tips at once and post-aspirates a trailing air gap that seats the plug above the tip orifice. The filled tips are positioned over a UV source beneath the deck and polymerized in place under a graded cure that drives conversion without overheating, with a brief final exposure of the ejected tip box to complete it. After a short bench rest and methanol wash, a full box of 384 tips is ready from a single run with ∼5 min of hands-on time (**Figure 1E, F**), two orders of magnitude faster than serial hand-packing. Because the Bravo aspirates a precise, identical volume of precursor into every tip and the cure proceeds in a thermally and optically controlled environment, the operator-dependent step of manual packing is replaced by an instrument-controlled one.

Scanning electron microscopy of a pSPEC plug extracted from the tip confirmed a well-defined monolith. At low magnification a cleaved plug appears as a continuous, self-supporting cylindrical body spanning the full tip lumen (**Figure 1G**), homogeneous in texture across the cross-section and without macroscopic voids, wall gaps, or channels along the polypropylene wall that could produce undesired preferential flow during centrifugation; the plug is a single integral piece of porous polymer rather than a packed bed of discrete particles. At high magnification (**Figure 1H**), the bulk material reveals the characteristic hierarchical morphology of a macroporous monolith: near-spherical primary particles of ∼100–300 nm are fused into globular aggregates of ∼1–2 µm, separated by an interconnected network of through-pores ∼0.5–2 µm wide. This architecture supplies both the open channels needed for low-backpressure centrifugal processing and the high surface area needed for capture, properties that can be fine-tuned by polymer composition, as we map next.

### The optimized monolith composition

We jointly optimized the chosen precursor for handling, sample preparation, and proteomic performance. In doing so we found that there is a narrow window in which a monolith tip functions at all for the purpose of a solid phase extraction material for proteomics sample preparation. An initial 96-parameter screen for polymerization showed this parameter region to be small, enclosed by three regions with distinct failure modes (**Figure 2A**): Firstly, above 31% total monomer percentage the plug becomes too dense to remain porous and no liquid can be centrifuged through it. Second, the crosslinker EDMA must reach at least ∼16% of the total solution otherwise there is too little difunctional crosslinker to form a stable network and the polymer never fully polymerizes. The third constraint was unexpected: because METAC carries the quaternary-ammonium groups responsible for anion exchange, we had assumed that more METAC would only improve binding; instead, while both low and high METAC fractions (as a percentage of total monomer) gave porous, workable tips, intermediate values polymerized into plugs that could not be used for sample preparation due to impermeability (U-shape, **Figure 2A**).

**Figure 2.**
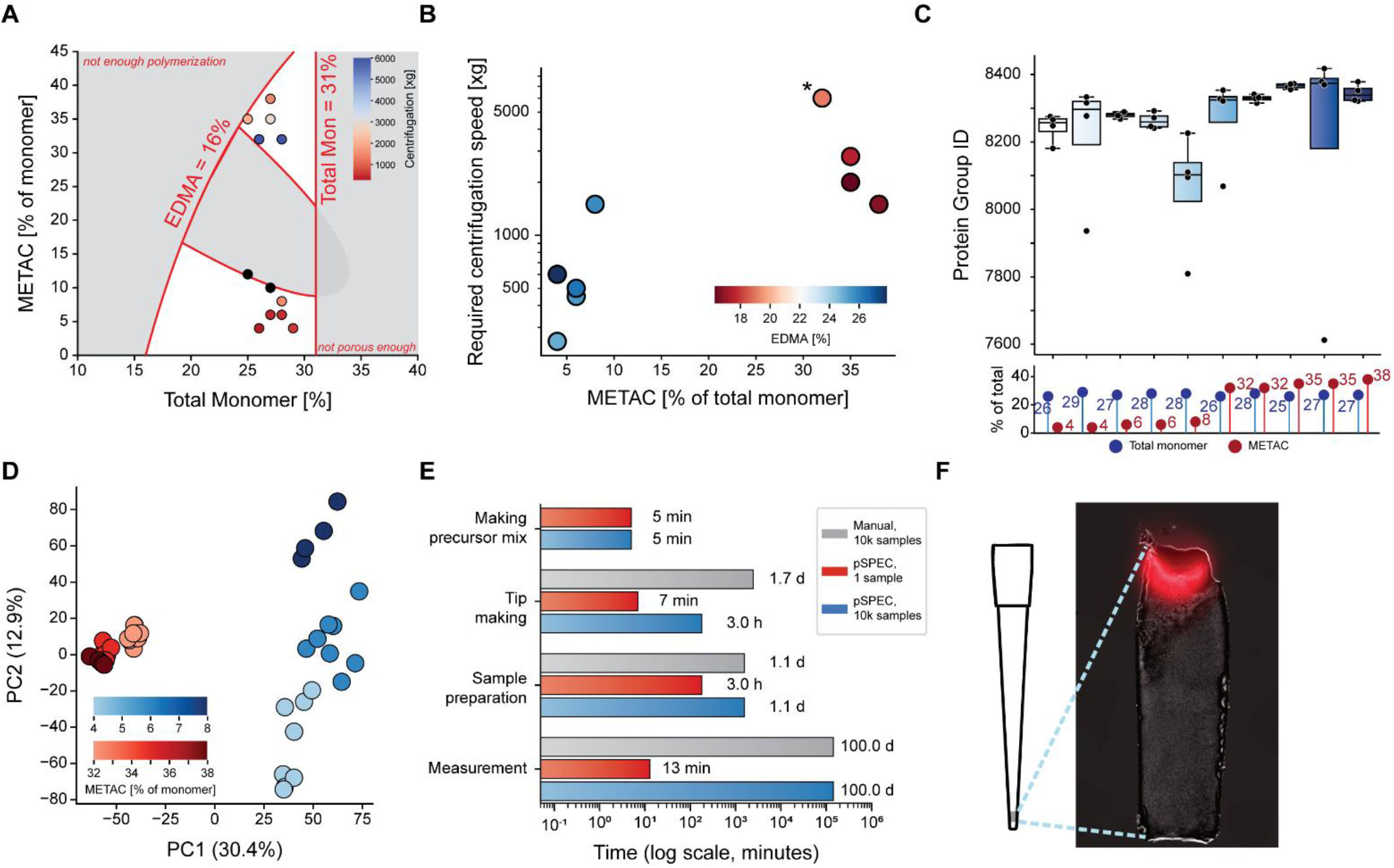
Design space mapping and selection of the optimized SAX monolith composition. (A) Polymer design space (total monomer % vs METAC % of monomer) overlaid with the three processability constraints: minimum crosslinker (EDMA = 16 %), maximum total monomer (= 31 %), and the lower ‘not porous enough’ boundary, which together delimit the viable window (grey region). Tested compositions are plotted and colored by the required centrifugal force (color bar, xg). (B) Backpressure per composition, expressed as the relative centrifugal force (xg) required to get 10 µL loading buffer through the tip in ∼90 s, versus METAC fraction and colored by EDMA content (mean, n = 4). The asterisk marks a composition whose value was extrapolated rather than directly measured. (C) Protein groups identified from 200 ng K562 lysate across the ten processable compositions (n = 4; box = IQR, center line = median, points = individual replicates). The composition of each tip is annotated below as total monomer % (blue) and METAC % of monomer (red). (D) PCA of log_10_protein-group intensities, colored by METAC fraction (separate scales for the low-METAC [4–8 %] and high-METAC [32–38 %] compositions). (E) Time per workflow stage (log_10_minutes) comparing manual sample preparation (grey; 10,000 samples), pSPEC at single-sample scale (red), and pSPEC at 10,000-sample scale (blue). (F) Alexa Fluor 750–labelled plasma proteins (250 ng) captured on a monolith tip and imaged in situ.

The usable region is thus constrained on three sides. Within it we screened 12 precursor mixes, seven in the low-METAC arm (25–29% total monomer, 4–12% METAC of total monomer) and five in the high-METAC arm (25–28% total monomer, 32–38% METAC of total monomer). The two mixes nearest to the impermeable band (27% monomer / 10% METAC and 25% monomer / 12% METAC) clogged irreversibly even under prolonged, high-speed spins. A clear physical rule emerged among the rest: the closer a composition was to either side of the middle band, the higher the resulting backpressure. We quantified this by measuring for each composition the centrifugal force needed to spin 10 µL of loading buffer through in ∼90 s (**Figure 2B**). This required 250 xg for the most permeable material and up to 6,000 xg near the U-shape. We found that within the low-METAC arm backpressure rose with increasing METAC, peaked sharply at the lower edge of the high-METAC arm, then fell again as METAC increased further, tracking the accompanying EDMA content. Therefore, the highest-METAC composition (27% total monomer, 38% METAC) cleared sample at a practical 1,500 xg, the lowest backpressure in the high-METAC arm, whereas other high-METAC conditions required ∼2,000– 2,500 xg for the same volume.

To characterize the performance gain of this composition at the proteome level, we processed 200 ng of K562 lysate on each of the ten compositions in quadruplicate. Identification depth was uniformly high (∼8,300 protein groups at 100 SPD) with a slight gain of 1.5% toward higher METAC (**Figure 2C**). This was a first hint that denser SAX functionality (greater percentage of METAC) is advantageous. The same ∼5% low-to-high-METAC increase appeared across more than 130,000 precursors, of which ∼70% were shared by every composition. The largest composition-specific set belonged to the high-METAC tips (**Supplementary Figure 1A, B**). The increased functionality therefore contributes genuine additional identifications. These differences were physicochemical, not stochastic: principal-component analysis (PCA) was dominated by the METAC fraction, which separated low from high METAC and ordered the high-METAC compositions monotonically along PC1 (∼30%), while PC2 (∼13%) resolved the gradations within the low-METAC arm (**Figure 2D**; the identical pattern recurred at precursor level, **Supplementary Figure 1C**). The polymer composition thus behaves as a proteome-level design variable, improving protein capture by physicochemical properties. Higher depth, more unique precursors, and, among the high-METAC tips, the lowest backpressure and easiest handling together led us to select 27% total monomer with 38% METAC (Mon27Met38) for the remainder of our study.

Once the composition is fixed, tips can be produced at scale without fabrication limiting throughput (**Figure 2E**). The preparation of the precursor mix takes about 5 min and is sufficient for 100 to 10,000 tips based on need. Tips are subsequently produced in 384-tip batches at 5-min production time. In this manner, making 10,000 tips takes no more than 3 h versus about 1.7 d to hand-pack the same number without any interruption (at 15 sec per tip for an experienced person). Downstream sample preparation steps amount to about 1 day from lysate to loaded Evotips for 10,000 samples, while measurement times are identical for packed and polymerized tips. With a 100 SPD LC-MS method, such a cohort would take 100 days to complete. What matters for a platform is therefore not the absolute duration of any single step but where the bottleneck is: in pSPEC the mix is made once and tips are produced by robot in minutes, so fabrication keeps pace with sample preparation at any realistic batch size and is not the limiting step. With hand-packed tips, fabrication alone would dominate the upstream workflow and make large batches impractical.

We next visually characterized how the polymerized monolith captures protein, since it was possible that the in situ material could behave differently from the packed Teflon-based material of our established SPEC workflow. Loading an Alexa Fluor 750–labelled plasma dilution series from 100 pg to 20 µg, the polymerized plug accepted up to ∼3–5 µg before clogging. (Note that if this happens, residual liquid can simply be withdrawn and the sample preparation workflow continued.) Fluorescence imaging of a 250 ng loaded tip shows the protein bound almost entirely in the uppermost ∼10% of the ∼500 nL plug (roughly 50 nL; **Figure 2F**). On-tip digestion therefore proceeds in an extremely small effective volume, the very confinement that makes SPEC efficient (**Figure 1A**).

Two final experiments defined the workflow’s practical limits and confirmed its robustness. The precursor combines polar and non-polar components and might phase-separate if mixed in an unfavorable order. We found that adding 1-propanol first as a bridging solvent prevents this. To show that such a phase separation does not propagate into the tips, we polymerized tips from a single batch at 10 timepoints across 0–24 h (six tips each) and processed plasma with every batch and measured each on the Orbitrap Astral. The proteome was invariant across the series with constant identifications (∼650 protein groups), ∼12% median CVs, and no temporal separation in PCA, differential testing, or correlation clustering (**Supplementary Figure 2**). This defines a working window of at least 24 h between mix-making and tip-making, effectively removing the timing of polymerization relative to mixing as a variable. Second, we optimized the loading-centrifugation speed using a demanding low-input sample, 5 ng of K562 lysate analysed on a timsTOF SCP. Identifications, CVs, and MS1 signal were essentially constant from 100 to 4,000 xg (**Supplementary Figure 3**). A marginal low-speed gain in data completeness (∼5,000 vs ∼4,720 protein groups across all six replicates, reflecting optimal binding rate physics from slow flow-through speed) was outweighed by the impractical ∼1.5 h spin at 100 xg, while every speed from 500 to 2,000 xg was indistinguishable. Based on these results, we adopted 1,000 xg loading in our pSPEC workflow.

In summary, we optimized conditions for in situ SAX-polymer formation in pipette tips for proteomics sample preparation by solid phase extraction capture in a high-throughput manner. Our trimethylammonium functionalized monolith captures protein material in the top 50 nL volume of porous polymer and can be handled in a minute timeframe at 1000 xg centrifugation speed. In this geometry up to 5 µg can be loaded; at these higher inputs the captured protein extends further down the monolith rather than remaining confined to its uppermost ∼10%.

### pSPEC delivers a deeper, faithful, and reproducible plasma proteome

Having mapped the polymer design space and benchmarked performance on K562 lysate, we tested the pSPEC workflow in more challenging conditions. We reasoned that plasma proteomics would benefit greatly from the high-throughput tip production, as hand-packing of tips for studies in the 1000s of samples is highly laborious, demands a very experienced operator, and is difficult to keep reproducible at that scale. As a complex biofluid, plasma also represents a stress test: its extreme dynamic range and small-molecule matrix arguably make it the most demanding routine proteomics matrix. Thus, a method that is deep, faithful, and reproducible for plasma should transfer to less demanding sample types. We therefore set out to benchmark pSPEC head-to-head against a standard in-solution neat-plasma workflow on the same LC-MS setup before using it to probe reproducibility.

Interestingly, our results indicated that pSPEC identified an additional 18% of proteins (mean of 620 vs 730 protein groups (in-solution vs pSPEC); **Figure 3A**) and 22% more precursors (mean of 5,400 vs 6,600; **Figure 3B**). This depth was additive: pSPEC recovered essentially all proteins also identified in the standard in-solution workflow while adding more than 150 protein groups (**Figure 3C**), extending the proteome rather than changing the covered proteome space physicochemically. Median CVs were only slightly higher for pSPEC (binning proteins into terciles by their in-solution-plasma signal, 5% vs 10% high tercile, 10% vs 15% mid tercile, 17% vs 22% for low tercile proteins; **Figure 3D**). pSPEC thus matches the reproducibility of the established workflow while enabling deeper plasma proteome measurement.

**Figure 3.**
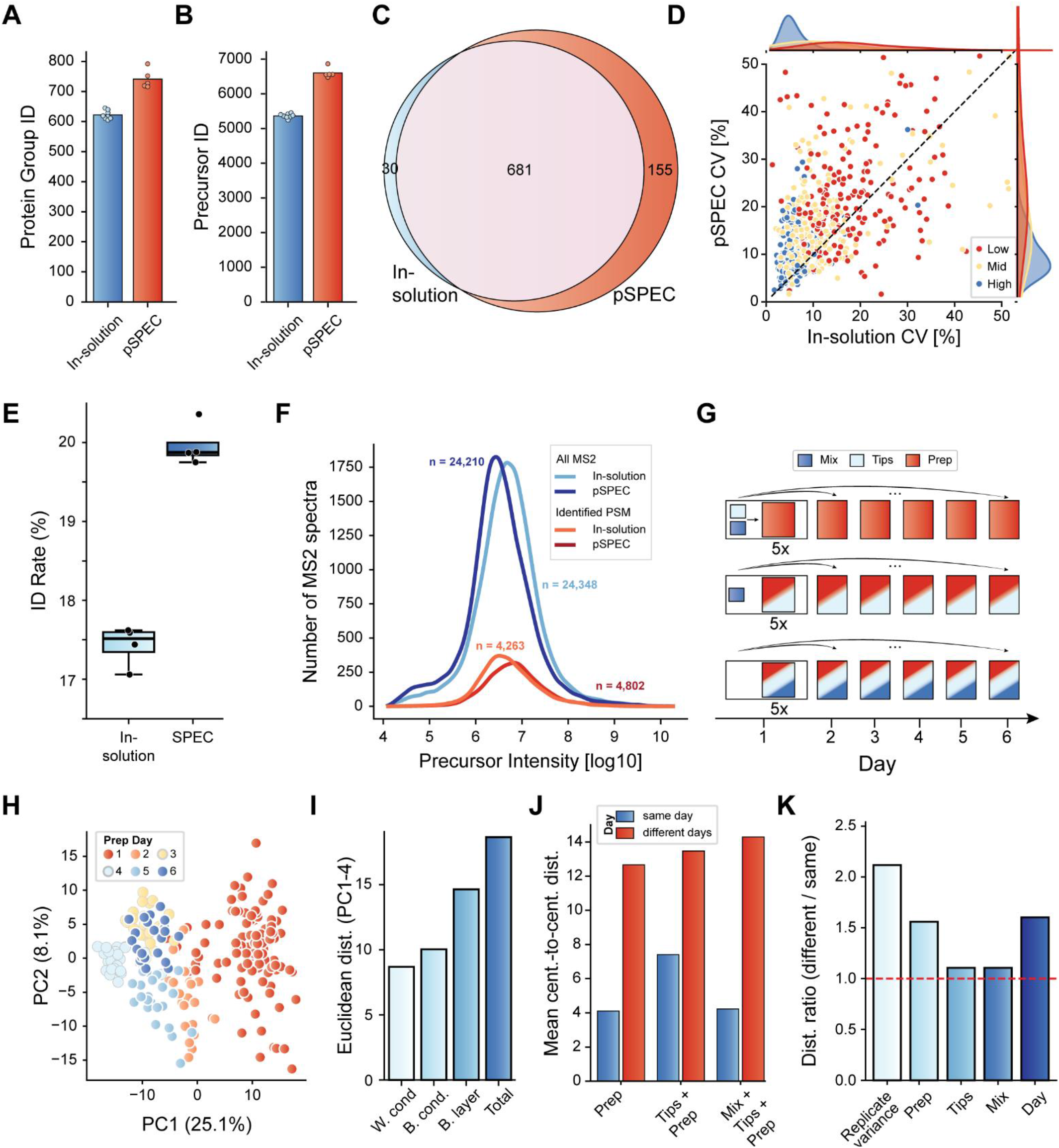
pSPEC plasma benchmark and reproducibility. (A, B) Protein groups (A) and precursors (B), neat in-solution plasma vs pSPEC (box = IQR, line = median, points = replicates). (C) Overlap of protein groups identified by in-solution plasma and pSPEC. (D) Median CV by in-solution-plasma abundance tercile (high/mid/low), in-solution vs pSPEC. (E, F) Matched DDA runs: MS2 scans (E) and identification rate (F), in-solution vs pSPEC. (G) Nested reproducibility design: Prep, Tips+Prep, Mix+Tips+Prep; 5× on Day 1 and 1× on each of Days 2–6, one pooled plasma, single LC-MS run. (H) PCA of log_10_protein-group intensities (top 500 proteins), colored by day. (I) Euclidean distances (PC1–4) by variability level (box = IQR, line = median). (J) Mean condition-centroid distances, same-day vs different-day. (K) Per-factor distance ratio.

We recently showed that SAX material such as magnetic beads preferentially capture cellular contaminants like platelets or PBMCs (*18*). To test whether our pSPEC workflow for plasma is affected by this problem, we conducted a serial dilution experiment by titrating platelets into plasma (100 to 2×10^6^ platelets/µL, plus a control without spike-in) (*19*), processed in parallel by the in-solution, pSPEC, and bead-based SAX workflows. Platelet load raised identifications for all three methods, but to very different degrees: at the highest spike-in, the SAX beads inflated protein groups several-fold to about 5,700, whereas pSPEC identified 2,940 proteins, closely tracking in-solution plasma (2,770 proteins) at every level (**Supplementary Figure 4A, B**). The platelet contamination index captures the same trends with near-identical scores for pSPEC and in-solution, but sharply elevated scores for the beads (**Supplementary Figure 4C**). The additional proteins pSPEC recovers are therefore bona fide plasma proteins, setting the monolith chemistry apart from bead-based SAX. We also speculated that pSPEC purification might lead to overall cleaner plasma samples, which would be an additional robustness advantage for clinical cohorts. Indeed, in matched DDA runs, in-solution and pSPEC generated essentially the same number of MS2 scans, yet pSPEC raised the identification rate from 17.4% to 20.0% (**Figure 3E, F**), reflecting fewer scans going to non-peptide or in general non-database search identifiable features per unit of acquisition.

Next, we turned to test reproducibility of polymer production and sample preparation with pSPEC as a part of the entire workflow. To this end, and inspired by clinical test architectures (*20*), we built a nested design that cumulatively varies the three steps that produce a sample ready for analysis: precursor-mix preparation (Mix), tip fabrication (Tips), and SPEC sample preparation (Prep) (**Figure 3G**). This design allows us to disentangle three layered potential sources of variance: Prep (reuse one mix and one tip batch; redo only the sample preparation), Tips + Prep (reuse the mix; fabricate a new tip batch and prep), and Mix + Tips+ Prep (everything made from scratch). We ran each condition five times on Day 1 (same-day variability) and once on each of Days 2–6 (different-day variability), all from a single plasma sample. All samples were acquired in single LC-MS runs so that instrument performance drift could not compromise this test of workflow variance.

Identification depth was constant across all 280 experimental runs (median of about 820 protein groups and 7,000 precursors; **Supplementary Figure 5A, B**). To reduce confounders from plasma’s low-abundance proteins, which carry intrinsically high CVs (**Supplementary Figure 5C)**, we restricted downstream analyses to the 500 most abundant proteins. Tip-to-tip CVs were then close to the baseline LC-MS variance floor (∼17– 22% protein, ∼22–28% precursor; **Supplementary Figure 5D, E**) compared to repeated injections of a single preparation with median CVs of 16% and 25%. PCA exposed the dominant structured source of variance: PC1 (25.1%) separated samples almost entirely by measurement day, with same-day samples clustering tightly regardless of which workflow components had been varied (**Figure 3H**). The separation was non-monotonic: each day formed its own cluster rather than drifting in sequence, pointing to discrete day-to-day batch effects, with PC2–PC4 contributing only minor additional structure (**Supplementary Figure 5G–I**).

Reducing each sample to its first four principal components (together 46.3% of the variance; Supplementary Figure 5F) and measuring Euclidean distances revealed the hierarchy of variance. As expected, distances grew with each additional workflow part, but the single largest variance was caused by the replication baseline, not any workflow layer (**Figure 3I**). Comparing condition centroids, which average out that noise, adding a new tip batch and a new mix on top of a new prep raised the different-day distance by only ∼6% (**Figure 3J**, with hierarchical clustering of these centroid distances in **Supplementary Figure 5J**). At the per-factor distance ratios, replicate variance was largest at 2.12, followed by day (1.60) and prep (1.56, itself largely inherited from the day effect, since preparations were nested within days), whereas tip fabrication and precursor mixing were close to the no-effect line (1.11 each; **Figure 3K**; absolute distances in **Supplementary Figure 5K**).

We conclude that the residual variance of the pSPEC plasma workflow is dominated by factors common to all proteomics such as the LC-MS measurement and day-to-day batch effects. Steps unique to pSPEC like precursor mix preparation and tip fabrication do not add additional variance. This establishes in situ polymerization as a controlled, reproducible production step. Practically, this argues for processing large cohorts in a single day and batch wherever possible, while defining tip manufacturing as a negligible reproducibility issue.

### pSPEC for body fluids

Encouraged by the plasma results, we next applied the identical workflow to three biofluids of very different protein content and concentration. We collected plasma, urine, and saliva from 18 healthy individuals (18 plasma, 14 urine, 16 saliva, see Methods), prepared every sample in triplicate, and acquired each by single-shot LC-MS (**Figure 4A**). This healthy cohort sets a deliberately stringent bar for urine: unlike proteinuria or diseased urine, healthy-donor urine is usually of low protein content. Thus, the depth reached here reflects the workflow and not an elevated protein input.

**Figure 4.**
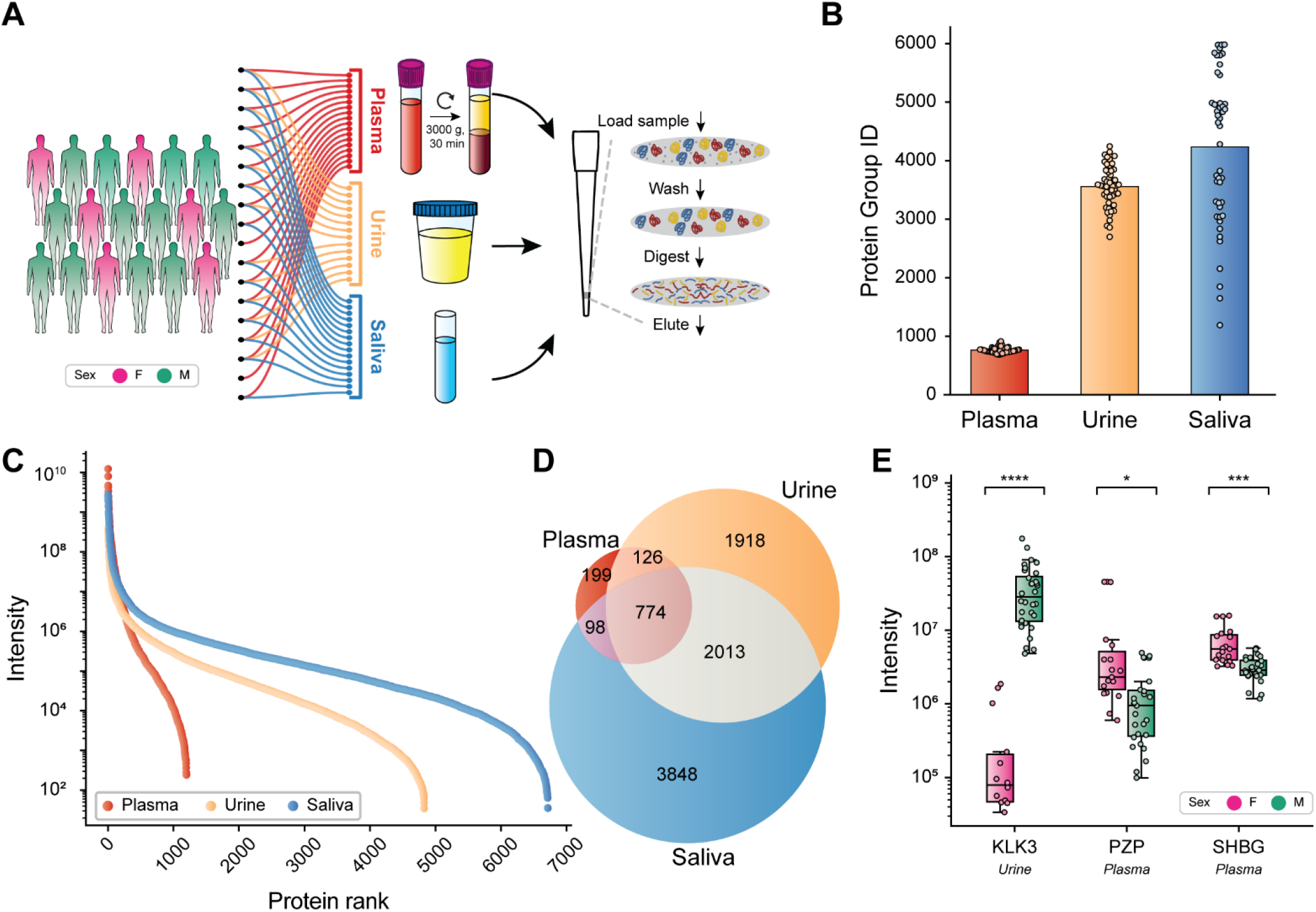
Deep, fluid-resolved plasma, urine, and saliva proteomes from a paired-donor cohort. (A) Cohort and workflow: plasma, urine, and saliva from 18 healthy donors (18 plasma, 14 urine, 16 saliva; not all donors provided all three fluids, 14 provided all three; sex coded female magenta / male green), each prepared in triplicate by the SPEC workflow and measured by single-shot LC–MS. Plasma was cleared at 3,000 xg for 30 min, urine and saliva at 3,000 xg for 20 min. (B) Protein groups per sample by fluid; bars show the mean and points are individual samples (triplicate per donor). (C) Rank-abundance per fluid: log_10_protein intensity versus protein rank (proteins ranked by descending mean intensity within each fluid). (D) Venn diagram of protein groups detected in each fluid. (E) log_10_intensity of sex-associated markers by donor sex (female magenta, male green; box = IQR, center line = median, points = individual replicates): KLK3 in urine, PZP and SHBG in plasma.

Across the three fluids pSPEC identified a median of ∼750 protein groups per plasma sample (∼1,050 across the cohort), a remarkable 3,500 per urine sample (∼5,000 total), and ∼4,800 per saliva sample (∼7,000 total), with ∼7,200, ∼22,700, and ∼32,000 precursors per sample, respectively (**Figure 4B, Supplementary Figure 6C**). These depths reflect two opposite challenges and how the monolith meets each. Plasma is capped by its intrinsic biology: a few proteins such as albumin and the immunoglobulins dominate the signal (**Figure 4C**), therefore no front-end method can deeply characterize plasma without depletion. In this context, pSPEC’s contribution is in-tip concentration and cleanliness rather than raw depth (**Figure 3**). For urine and saliva MS-based proteomics is primarily limited by protein concentration rather than dynamic range alone. pSPEC’s confinement is an ideal match for this challenge: all proteins of a sample are captured and concentrated on the SAX material into a ∼50 nL digestion volume regardless of starting volume, keeping on-tip digestion extremely efficient (**Figure 1B**).

For urine of healthy donors, single-injection untargeted analyses have typically reported ∼1,000–1,500 protein groups (∼1,140 by an automated HILIC-bead workflow and fewer in earlier single-run studies) with only the most recent, optimized DIA methods reaching ∼3,000 in long LC gradients and deeper catalogues requiring extensive fractionation across pooled donors (*21*– *23*). pSPEC reaches ∼3,500 in a 12.95 min LC-MS run. This principle, concentrating a dilute proteome onto a small bed for efficient digestion, has been leveraged to some extent by pioneering bead- and membrane-based methods such as HILIC-bead and MStern protocols (*4, 21*). pSPEC pushes it further, confining digestion to a minimal reaction volume while stringently washing away non-protein sample content, and without the lengthy bead equilibration and incubation those protocols require. Depth in urine, however, scales with the cohort: ageing and disease raise urinary protein excretion, so recent single-shot studies of older or diseased populations report comparable or higher counts, for instance a median of ∼3,800 proteins per individual in a Parkinson’s cohort (*24*). Our donors, by contrast, were healthy volunteers mostly aged 20–35, in whom urine is intrinsically protein-poor, so the ∼3,500 protein groups reached here reflect genuine analytical depth rather than disease- or age-driven proteinuria. Saliva, where single runs commonly yield ∼1,000– 2,000 human protein groups (*25*–*27*), shows the same advantage with an average of ∼4,800 protein groups identified across individuals. Quantification was well reproducible across all three fluids: per-individual CVs across triplicate stayed below ∼15% (plasma), ∼22% (urine), and ∼17% (saliva) at the protein-group level (**Supplementary Figure 6A, B, F**).

Each fluid contributed a large specific proteome: 3,848 protein groups unique to saliva, 1,918 to urine, and 199 to plasma, with only 774 shared by all three (**Figure 4D**). Interestingly, urine and saliva resemble each other more than either resembled plasma in Spearman correlation clustering (**Supplementary Figure 6E, G**). A curated panel of literature-defined marker proteins was elevated specifically in its source fluid: ALB, FGA, FGG, APOA1, APOB, and C3 in plasma (*28, 29*); the kidney and urinary-tract proteins UMOD, CUBN, LRP2, EGF, and FABP1 in urine (*28*–*30*); and the salivary proteins AMY1A, CST2, CST5, PIP, and CA6 in saliva (*27*) (**Supplementary Figure 6K**).

Finally, the workflow recovered established sex biology although this was not a dominant driver of variance within any fluid (Supplementary Figure 6H–J); a Welch’s t-test between male and female donors still yielded the expected sex-specific proteins as the dominant hits (Supplementary Figure 6L–N). The clearest ones are textbook markers in the expected fluids and directions (**Figure 4E**): the prostate-derived kallikrein KLK3 (PSA) was on average 250-fold more abundant in male urine (p < 0.0001), while SHBG (p < 0.001) and PZP (p < 0.05) were more abundant in female plasma.

## Discussion

SPEC draws its depth at low input from confinement: capture, washing, on-tip digestion, and elution all proceed within the nanoliter pore volume of the stationary phase, without depletion or fractionation (*12*). pSPEC retains that mechanism and changes only how the phase is built, replacing hand-packed beads or discs with a monolith photopolymerized in place from a defined recipe. Casting the bed by light rather than assembling it mechanically has two consequences. It can be produced at scale and at low cost: 384 tips from a single robot run in minutes, from inexpensive bulk monomers. And because curing requires only optical access, not a fixed packing geometry, the same chemistry can be formed in almost any volume, whether shrunk toward the low or sub-nanoliter beds that single-cell inputs require, where dead volume and surface losses dominate, or enlarged to the capacity needed to incorporate an affinity-capture step into the confined workflow (*31*). Producing the SPEC phase economically, at scale, and in a geometry matched to the application is the principal advance reported here.

Creating tips in quantity is useful only if they are interchangeable, and the nested reproducibility design shows that they are. Renewing, cumulatively, the precursor mix, the tip batch, and the sample preparation, the steps unique to in situ polymerization showed essentially no effect: a fresh mix and a new tip batch each changed inter-day distances by only a few percent, while the predominant sources of variance were the LC-MS measurement and the day-to-day batch effects that are inherent to proteomic workflows (*32*). In situ polymerization is thus a controlled production step rather than a source of error to be managed, which in practice argues for preparing large sample sets in a single batch and on a single day.

Defining the phase by a disclosed recipe also makes its properties adjustable: charge density, porosity, and capacity are set by composition, and we mapped the narrow window in which a functional monolith forms together with its three failure boundaries (*16*). The composition is therefore open for others to reproduce and to push toward separation chemistries beyond the single SAX formulation used here.

The second result of broad importance is the depth reached in dilute body fluids. With one chemistry, one load-wash-digest-elute protocol, one MS method, our workflow returned 3,500 protein groups from a single short injection of healthy urine and 4,800 from saliva, depths that have generally required depletion, fractionation, or laborious and potentially proteome-distorting bead protocols (*21*–*23, 25*–*27, 33, 34*). The mechanism is intrinsic to SPEC rather than to the monolith: a dilute proteome presented in a large aqueous volume is concentrated onto the ∼50 nL bed and digested at high effective concentration. What pSPEC adds is the ability to deliver that advantage reproducibly and at scale, across fluids, from a single workflow. Plasma, by contrast, is bounded by its own dynamic range rather than by the tip, consistent with an enrichment that extends the measurable proteome without reshaping it.

These depths reflect fluid-appropriate biology, not artifact. In plasma, a SAX phase invites the objection that the extra identifications are enriched cellular contamination; a platelet titration ruled this out, with pSPEC tracking in-solution neat plasma across the whole spike-in range while bead-based SAX inflated identifications several-fold by capturing the contaminating material. The samples were also measurably cleaner, raising the identification rate from 17% to 20% at matched acquisition. Cleaner input of this kind should keep the instrument clean for longer and may make FAIMS unnecessary, both practical advantages at cohort scale. The workflow recovered textbook sex-dimorphic markers in the correct fluids and directions: KLK3 in male urine, SHBG and PZP in female plasma (*35*). The added depth therefore converts into interpretable signal rather than a longer list of marginal hits.

Cast from a robust recipe and formed by light, these tips can be produced economically, at scale, and in the geometry an application calls for, while remaining reproducible enough to be treated as a settled part of the workflow. We believe that pSPEC will become a routine and universal part of many proteomics workflows.

## Materials and Methods

### Human samples

Human samples. All human biofluids used in this study, plasma, urine, and saliva, were obtained from healthy adult donors after written informed consent. The study was approved by the Ethics Committee of the Max Planck Society for the Advancement of Science (Reg. No. 2025_35), and all experiments conformed to the principles of the WMA Declaration of Helsinki and the US Department of Health and Human Services Belmont Report.

### Blood collection and plasma preparation

Whole blood was collected by venipuncture into EDTA-containing tubes (9 mL). Immediately after collection, tubes were gently inverted 10 times to ensure anticoagulant mixing, followed by centrifugation at 3,000 xg for 30 min. Only the upper two-thirds of the plasma layer was carefully aspirated from above the buffy coat, leaving the lowermost plasma and the cellular interface undisturbed to minimize platelet and cell carry-over, and stored at −80 °C until further processing.

### Isolation of purified platelets

For experiments requiring defined platelet contamination levels, blood components were fractionated by differential centrifugation. Whole blood was first centrifuged at 500 xg for 7 min to separate platelet-rich plasma from cellular components. The supernatant (platelet-rich plasma) was carefully transferred to a new tube and centrifuged again at 500 xg for 7 min. This supernatant was collected and centrifuged at 3,000 xg for 7 min. For platelet isolation, the pellet from the 3,000 xg centrifugation was resuspended in 4 mL PBS/EDTA (1.6 mg/mL EDTA), centrifuged at 3,000 xg for 7 min, and the supernatant was discarded. This washing step was repeated once more, resulting in purified platelets.

### Urine collection and preparation

Urine was collected by donors into a sterile collection cup, cleared of cellular debris by centrifugation at 3,000 xg for 20 min at 4 °C, and the supernatant transferred and stored at −80 °C until further processing.

### Saliva collection and preparation

Saliva was collected by passive drool into a tube, cleared of cellular debris by centrifugation at 3,000 xg for 20 min at 4 °C and immediately frozen at −80 °C without further processing.

### Monolith precursor preparation

Precursor solutions for SAX monolith polymerization were prepared by combining the functional monomer [2- (methacryloyloxy)ethyl]trimethylammonium chloride (METAC, 75% aqueous solution; Merck), the crosslinker ethylene glycol dimethacrylate (EDMA; Merck), and a ternary porogen system of 1-propanol, hexanol, and water (nominal porogen composition 52:33:15 by volume of the directly added porogen components). Polymerization was initiated photochemically using 2,2-dimethoxy-2-phenylacetophenone (DMPA; Merck), added as a 10% (w/v) stock solution in 1-propanol and constituting 10% of the total precursor volume. All components were combined in a 250 mL glass bottle in the order: 1-propanol, METAC, hexanol, EDMA, water, and DMPA stock. After addition of each component, the mixture was thoroughly vortexed to prevent phase separation.

A representative 210 mL batch of the high-SAX precursor (27% total monomer, 38% METAC of total monomer volume) contained 68.8 mL 1-propanol, 21.5 mL METAC (75% aqueous), 43.7 mL hexanol, 35.2 mL EDMA, 19.8 mL water and 21.0 mL DMPA stock (10% w/v in 1-propanol), and prepared to a total volume of 210 mL. Batch volume was scaled as required.

### Automated tip fabrication

Monolith tips were fabricated by UV-initiated in situ polymerization inside standard 30 µL polypropylene pipette tips using an Agilent Bravo automated liquid handling platform. The precursor solution was transferred into a reservoir (trough) with a liquid column height of approximately 3.5-4 cm and allowed to settle for 1 h at room temperature in the dark.

The Bravo aspirated the precursor solution into 384 pipette tips simultaneously (one full tip box per run) in approximately 5 min of active instrument time. Polymerization was performed in three stages directly following aspiration. In the first stage, tips remained on the Bravo head and were exposed to UV light (Hoenle, LED Cube 100 IC, 365 nm) at 20% power for 3 min to gently initiate polymerization and begin gel formation without overheating. In the second stage, UV power was increased to 100% for 1 min to drive polymerization to higher conversion and solidify the monolith structure. For the third stage, tips were ejected back into the tip box, which was removed from the Bravo and exposed to UV light in a UV chamber at 100% power for 1 min to ensure complete curing throughout the monolith plug.

After polymerization, tips were allowed to cool and dry for approximately 1 h at room temperature. Tips were then washed once with 30 µL methanol (MeOH) to remove residual porogen, unreacted monomers, sol fraction oligomers, and photoinitiator fragments. After washing, tips were ready for use in the SPEC workflow or stored at room temperature until use.

### Polymer characterization

For scanning electron microscopy (SEM), the dried polymer plugs were extracted from pipette tips by retrograde probing and a light push using a Pt-Ir (90:10 w/w) wire (0.25 mm diameter, Thermo Fisher Scientific). The pipette tip was then turned upside down and tapped on a cleaned object slide to release the resin plug. A carbon conductive sticky tab (Ted Pella) was mounted on an SEM sample stub (Ted Pella). The polymer plug was transferred onto the adhesive carbon surface under a dissecting stereo microscope with the help of an eyebrow hair mounted on a wooden stick. The plug was split in half on the adhesive film using a new scalpel blade. The halves were placed side by side by lightly pushing outwards on the freshly dissected face with the scalpel blade flanks. The sample was mounted on a stub holder shuttle and transferred into a FIB/SEM instrument (Aquilos 2, Thermo Fisher Scientific). The sample was rendered conductive deploying the integrated Pt sputter coater (0.10 mbar, 30 mA, 15 s). Imaging was performed at an acceleration voltage of 3 kV and 50 pA beam current at different magnifications, as indicated by the scale bars in the micrographs shown.

### Visualization of the tip

Plasma labeling with Alexa Fluor 750 For visualization of plasma binding on the monolith tip, plasma proteins were fluorescently labeled using the amine-reactive NHS ester of Alexa Fluor 750 (Thermo Fisher Scientific, Cat. No. A20011). Plasma was combined with the dye and incubated for 1 h at room temperature in the dark to allow conjugation of the dye to primary amines on plasma proteins. Unreacted dye was removed by passing the reaction through a Zeba Spin desalting column (7 kDa MWCO, 0.5 mL format; Thermo Fisher Scientific, Cat. No. 89882) according to the manufacturer’s protocol. The labeled plasma was then processed by the standard SPEC workflow on the monolith tip.

#### Imaging

For fluorescence microscopy, the plugs were removed from the pipette tips as for scanning electron microscopy. Glass slides were coated with CoverGrip Coverslip Sealant (Biotium, #23005) and the adhesive was allowed to dry down lightly. Then, the plug was maneuvered onto the adhesive layer and subsequently split as for the SEM preparation. Slides were imaged on a Zeiss Axioscan 7 (ZEN blue) with a Plan-Apochromat 20×/0.8 M27 objective. Acquisitions were 9-plane z-stacks (0.81 µm step, 6.48 µm total) recorded with 2×2 binning on an Axiocam 712 monochrome camera at 14-bit depth. Two channels were collected per tile: a transmitted-light brightfield channel (TL LED lamp, 300% intensity, 10 µs flash; acquired for TIE focus per the scan profile) and a far-red fluorescence channel for the Alexa Fluor 750 protein label (AF750; Colibri 7 735 nm LED module at 10% intensity, excitation/emission maxima 752/779 nm, captured through the 720-750 nm excitation and 770-800 nm emission bands of the 112 HE multiband filter set). Tiles were stitched in ZEN Blue and stitched scenes exported as TIFF files.

### Sample preparation workflows

#### K562 lysate

MS-compatible human protein extract from K562 cells (Promega, Cat. No. V6941, 1 mg) was used as a standardized protein source. The lysate was diluted to 1 mg/mL in 50 mM triethylammonium bicarbonate (TEAB) pH 8.5, 10 mM TCEP, 40 mM chloroacetamide (CAA) and incubated at room temperature for 1 h. Prior to loading, the lysate was adjusted to the required concentration by dilution in 50 mM TEAB, 0.01 % N-Dodecyl β-D-maltoside (DDM) (0.04 mg/mL for 200 ng).

#### Plasma

Plasma protein concentration was assumed to be 50 µg/µL. 1 µL of plasma was combined with 24 µL of buffer (5% sodium deoxycholate (SDC), 0.01% DDM, 10 mM Tris(2-carboxyethyl)phosphine (TCEP), 40 mM CAA in 100 mM Tris-HCl, pH 8.5), giving a nominal concentration of 2 µg/µL. The sample was further diluted 1:40 in the same buffer to a final concentration of 50 ng/µL and heated at 95 °C with agitation for 10 min on a Thermomixer C (Eppendorf). After cooling to room temperature, samples were processed by the SPEC workflow.

#### Urine

40 µL of cleared urine was supplemented with TCEP and CAA to final concentrations of 10 mM and 40 mM, respectively, and incubated at room temperature for 30 min. Because of urine’s low protein content, and the resulting need for high input volumes, the SPEC workflow for urine was performed by adding the reduced and alkylated urine to an equal volume of 40 mM CAPS (pH 10.5), 0.02 % DDM, 90% MeOH, in place of the standard loading buffer. Because total protein concentration in urine varies substantially between donors, peptide concentration was measured after the SPEC workflow by A280 on a NanoDrop One (Thermo Fisher, Madison, USA) (1 Abs = 1 mg/mL peptide) and the volume corresponding to 400 ng was loaded onto an Evotip.

#### Saliva

Saliva protein concentration was assumed to be 1 µg/µL. 1 µL of saliva was combined with 19 µL of buffer (composition as for plasma) to a final concentration of 50 ng/µL and heated at 95 °C with agitation for 10 min on a Thermomixer C (Eppendorf). Samples were then processed by the SPEC workflow.

#### SPEC workflow using monolith tips

Samples were processed using the Solid-Phase Extraction Capture (SPEC) workflow (*12*) in a two-tip configuration, where proteins are captured on the SAX monolith tip, enzymatically digested within the confined pore space, and eluted as peptides onto a C18 Evotip for LC-MS analysis.

SAX monolith tips were first primed with 10 µL dimethyl sulfoxide (DMSO) (2000 xg, 3 min) and equilibrated with 20 µL equilibration buffer (20 mM CAPS, 0.01% DDM in 50% MeOH, pH 10.5; 2000 xg, 2 min). Protein lysate (up to 500 ng in up to 30 µL loading buffer consisting of 20 mM CAPS with 0.01% DDM in 50% MeOH) was loaded onto the SAX tip by centrifugation (1000 xg, 10 min). Tips were washed with 20 µL washing buffer (50 mM TEAB, 0.01% DDM; 2000 xg, 3 min). For on-tip digestion, 2 µL freshly prepared digestion buffer was added to each tip. The digestion buffer contained 50 mM TEAB, 10 mM TCEP, 0.01% DDM, and proteases (0.05 µg/µL each of trypsin and LysC). After a brief pulse spin (200 xg, 30 s), tips were incubated at 37 °C for 1 h.

For peptide recovery, Evotips (Evotip Pure, Evosep) were prepared by washing with 50 µL Evo B (700 xg, 1 min), soaking in isopropanol for 1 min, washing with 50 µL Evo A (700 xg, 1 min), and adding 100 µL Evo A followed by a pulse spin. The SAX monolith tip was placed above the prepared Evotip, and peptides were eluted with 20 µL elution buffer (1% formic acid, 0.01% DDM; 1000 xg, 2 min) directly onto the C18 material. The Evotip was then washed with 50 µL Evo A (700 xg, 1 min), and 100 µL Evo A was added with a final pulse spin before LC-MS analysis.

#### In-solution neat plasma workflow

Plasma (1 µL) was combined with 24 µL buffer (100 mM Tris pH 8.0, 40 mM chloroacetamide, 10 mM TCEP). Samples were heated at 95 °C with agitation for 10 min on a Thermomixer C. After cooling to room temperature, 10 µL digestion buffer (8 µL lysis buffer, 1 µL trypsin [0.5 µg/µL], 1 µL LysC [0.5 µg/µL]) was added, and proteins were digested at 37 °C for 16 h. Peptides were acidified and loaded onto Evotips according to the manufacturer’s protocol for LC-MS analysis.

### Data acquisition by mass spectrometry

Samples were analyzed on two LC-MS platforms: an Evosep Eno liquid chromatography system (Evosep) (*11*) coupled to either an Orbitrap Astral mass spectrometer (*36, 37*) (Thermo Fisher Scientific) or a timsUltra 2 mass spectrometer (Bruker Daltonics). Mobile phases consisted of 0.1% formic acid in water (buffer A) and 0.1% formic acid in acetonitrile (buffer B), and peptides were loaded onto C18 trap tips (Evotip Pure, Evosep) according to the manufacturer’s protocol prior to LC-MS analysis. Quality control samples were analyzed regularly throughout each analytical sequence to monitor system performance and stability. Three Orbitrap Astral acquisition methods, one timsTOF SCP method, and one Orbitrap Astral DDA method were used as detailed below.

*Orbitrap Astral, data-independent acquisition*. For plasma and K562 samples processed by SPEC, peptides were separated on an 8 cm Aurora Rapid XT UHPLC column (AUR4-80150C18-XT, Ionopticks) at 50 °C using the Evosep ‘100 samples per day’ method with pre-formed gradients and a total runtime of 12.95 min per sample; 200-500 ng of peptides were loaded per Evotip. For low-input 5 ng K562 samples, peptides were separated on a 5 cm Aurora Rapid XT UHPLC column (AUR4-50150C18, Ionopticks) at 50 °C using the Evosep ‘Whisper 80 SPD’ method with a total runtime of 16.3 min per sample. The ion source was operated with a static spray voltage of 1,900 V in positive ion mode and an ion transfer tube temperature of 280 °C. FAIMS (high-field asymmetric waveform ion mobility spectrometry) was used in standard resolution mode with a compensation voltage of −40 V and a carrier gas flow of 3.5 L/min. For plasma, urine and saliva samples, full MS1 scans were acquired in the Orbitrap analyzer at a resolution of 120,000 FWHM over 380–980 m/z (RF lens 40%, normalized AGC target 500%, maximum injection time 3 ms), and DIA MS2 scans covered the same mass range divided into 150 isolation windows of 4 Th with window placement optimization enabled, acquired in the Astral analyzer with an HCD collision energy of 25%, normalized AGC target of 500%, maximum injection time of 7 ms, and a fragment scan range of 150–2,000 m/z (*38*). For 200 ng K562 samples, the DIA scheme was tightened to 200 isolation windows of 3 Th over the same precursor range, with the MS1 maximum injection time set to 3 ms and the MS2 maximum injection time set to 6 ms. For 5 ng K562 samples on the Whisper 80 SPD gradient, MS1 resolution was raised to 240,000 FWHM with a maximum injection time of 100 ms, and DIA MS2 scans used 75 isolation windows of 8 Th with a maximum injection time of 14 ms; all other parameters were as for the plasma method. Data were collected in profile mode for MS1 and centroid mode for MS2.

*Orbitrap Astral, data-dependent acquisition*. For plasma DDA experiments, samples were separated under the same conditions as the plasma DIA method (Evosep 100 SPD, 8 cm Aurora Rapid XT, 50 °C, 250 ng load). FAIMS was used in standard resolution mode with a compensation voltage of −40 V and a carrier gas flow of 3.5 L/min, as for the DIA method. Full MS1 scans were acquired in the Orbitrap analyzer at a resolution of 120,000 FWHM over 350–1,050 m/z (RF lens 40%, normalized AGC target 500%, maximum injection time 7 ms), with monoisotopic peak determination set to ‘peptide’ and charge states 2–6 selected for fragmentation. Data-dependent MS2 scans were acquired in the Astral analyzer with a cycle time of 0.6 s, an isolation window of 1.4 Th, an HCD collision energy of 30%, a normalized AGC target of 1,000% (maximum injection time 20 ms), and a scan range of 100– 2,000 m/z. Dynamic exclusion was applied for 5 s with a mass tolerance of ±10 ppm, and an intensity threshold of 5 × 10^3^ was used for precursor selection.

*timsUltra AIP, dia-PASEF*. For low-input 5 ng K562 samples on the timsUltra AIP, peptides were separated on a 15 cm PepSep C18 column (1893474, 15 cm x 150 µm x 1.5 µm, Bruker) using the Evosep ‘Whisper 40 SPD’ method. The ion source was operated in positive mode with a capillary voltage of 4,500 V, dry gas flow of 8.0 L/min, dry temperature of 200 °C, and a nebulizer pressure of 0.3 bar. TIMS was operated over a 1/K_0_range of 0.70–1.30 V·s/cm^2^with a ramp and accumulation time of 100 ms each (duty cycle ∼100%). dia-PASEF MS scans covered a mass range of 100–1,700 m/z, with isolation windows and ramp design generated by py_diAID (*39*) to optimize precursor coverage across the m/z–ion mobility space.

### Data analysis / spectral search

The raw files were processed using DIA-NN (version 2.2.0) (*40*) on a high-performance computing cluster. A spectral library was predicted *in silico* with the DIA-NN deep-learning predictor from a human UniProt Swiss-Prot isoform database (UP000005640, 42,439 entries). Trypsin/P was specified as the protease with up to one missed cleavage, and the predicted library spanned peptide lengths of 7–30 residues, precursor charges of 1–4, precursor m/z of 300– 1,800, and fragment m/z of 200–1,800. Carbamidomethylation of cysteine was set as a fixed modification, and methionine oxidation and protein N-terminal acetylation as variable modifications (maximum one variable modification per peptide). All files were analysed with match-between-runs enabled (“--use-quant” and “-- reanalyse”), with peak centering, smart profiling, retention-time profiling, and relaxed protein inference. Mass accuracy was set to 10 ppm for MS1 and MS2 and the scan window to 7, and cross-run normalization used the default RT-dependent strategy. Protein quantification used MS1 and MS2 with no interference signal removal, and the false-discovery rate was controlled at 1% at the precursor level. Quantitative matrices were generated for downstream statistical analyses.

DDA raw files were processed with FragPipe (version 23.1) (*41*) using MSFragger as search engine, against the same human UniProt Swiss-Prot isoform database. Default parameters were used throughout: strictly tryptic digestion with up to two missed cleavages, peptide length 7–50, precursor and fragment mass tolerances of ±20 ppm with automated mass calibration, carbamidomethylation of cysteine as a fixed modification, and oxidation of methionine and protein N-terminal acetylation as variable modifications (maximum three per peptide). The false-discovery rate was controlled at 1% at the PSM, peptide, and protein level by Philosopher, and label-free MS1 quantification with match-between-runs was performed by IonQuant.

### Bioinformatics analysis

Downstream analysis was performed in Python within Jupyter notebooks, using pandas and NumPy for handling of the proteomics data matrices, and SciPy, matplotlib, and seaborn for statistics and visualization. Protein-level reports from DIA-NN were taken as input, and protein intensities were log_10_-transformed before any further processing. When a complete data matrix was required, for example as input to principal component analysis, missing values were filled in by K-nearest-neighbors imputation with n_neighbors = 5. Prior to downstream analysis, the DIA-NN output was filtered at a 1% false discovery rate by requiring Q.Value ≤ 0.01, Lib.Q.Value ≤ 0.01, and Lib.PG.Q.Value ≤ 0.01; only entries passing all three criteria were retained.

#### Outlier filtering

To retain the most consistent replicates per condition, we applied a consensus outlier-detection procedure that combines three complementary approaches at both the protein-group and the precursor level, all operating on the log_10_-transformed intensities. The correlation-based approach flagged, within each condition, the two replicates with the lowest median pairwise Pearson correlation to the remaining replicates. The PCA-based approach performed a global PCA on KNN-imputed (k = 5), z-scored intensities and flagged the two replicates with the greatest Euclidean distance from their group centroid in PC1–PC4 space. The perturbation-based approach exhaustively evaluated all C(n, 2) pairwise removal combinations within each condition and identified the pair whose removal minimized the median per-feature coefficient of variation across the remaining replicates; unlike the other two methods, this captures joint effects in which replicates appear acceptable individually but jointly inflate variance. Each of the three methods was applied at both data levels, yielding six flags per replicate. In the default combined-consensus mode, the two replicates per condition with the most flags were removed, with ties broken by the lowest median protein-group correlation.

Euclidean distance in PCA space. Distances between samples and between condition centroids in PCA space were computed across the first four principal components. For two points P and Q with coordinates (PC1, PC2, PC3, PC4), the distance is d(P, Q) = √[(PC1_P − PC1_Q)^2^+ (PC2_P − PC2_Q)^2^+ (PC3_P − PC3_Q)^2^+ (PC4_P − PC4_Q)^2^].

Platelet carry-over was quantified per sample using the platelet quality-marker panel of Geyer et al. (*19*): the index is the summed intensity of the detected platelet-marker proteins divided by the summed intensity of all quantified proteins, with higher values indicating greater platelet contribution. Indices were computed from DIA-NN protein-group intensities and compared across the in-solution, pSPEC, and bead-based SAX workflows.

Group comparisons (e.g. male versus female donors; aged versus freshly mixed precursor) were performed on log_10_-transformed intensities using a two-sided Welch’s t-test. Multiple-testing correction was applied by [Benjamini–Hochberg / permutation-based FDR], and proteins were considered significant at [adjusted p < 0.05 / p < 0.05 and |log_2_fold-change| > 0.5].

## Acknowledgments

We thank all members of the Proteomics and Signal Transduction Group for help and discussions. We thank Daniel Bollschweiler and Tillman Schaefer from the Cryo-EM core facility at the Max Planck Institute of Biochemistry for their assistance with electron microscopy imaging, and Igor Paron and Katharina Zettl for technical assistance. We thank Jörg Reichart for clinical assistance. We thank all body fluid donors who made this research possible. This project was supported by the Max Planck Society for the Advancement of Science.

## Author contributions

K.K.: Conceptualization, data curation, formal analysis, investigation, methodology, validation, visualization, writing - original draft, and writing - review & editing. L.H.: Data curation, formal analysis, visualization, and writing - review & editing. D.O.: formal analysis, visualization, and writing - review & editing. N.E.: formal analysis, visualization, and writing - review & editing. A.M.S.: Data curation, formal analysis, visualization, and writing - review & editing. T.H.: Conceptualization, methodology, and writing - review & editing. A.S.S.: formal analysis, visualization, and writing - review & editing. V.A.: formal analysis, visualization, and writing - review & editing. C.J.O.K.: Conceptualization, data curation, methodology, and writing - review & editing. M.M.: Conceptualization, funding acquisition, project administration, resources, supervision, writing - original draft, and writing - review & editing. J.B.M.R.: Conceptualization, data curation, formal analysis, investigation, methodology, resources, software, supervision, validation, visualization, writing - original draft, and writing - review & editing.

## Supplementary Figures

**Supplementary Figure 1.**
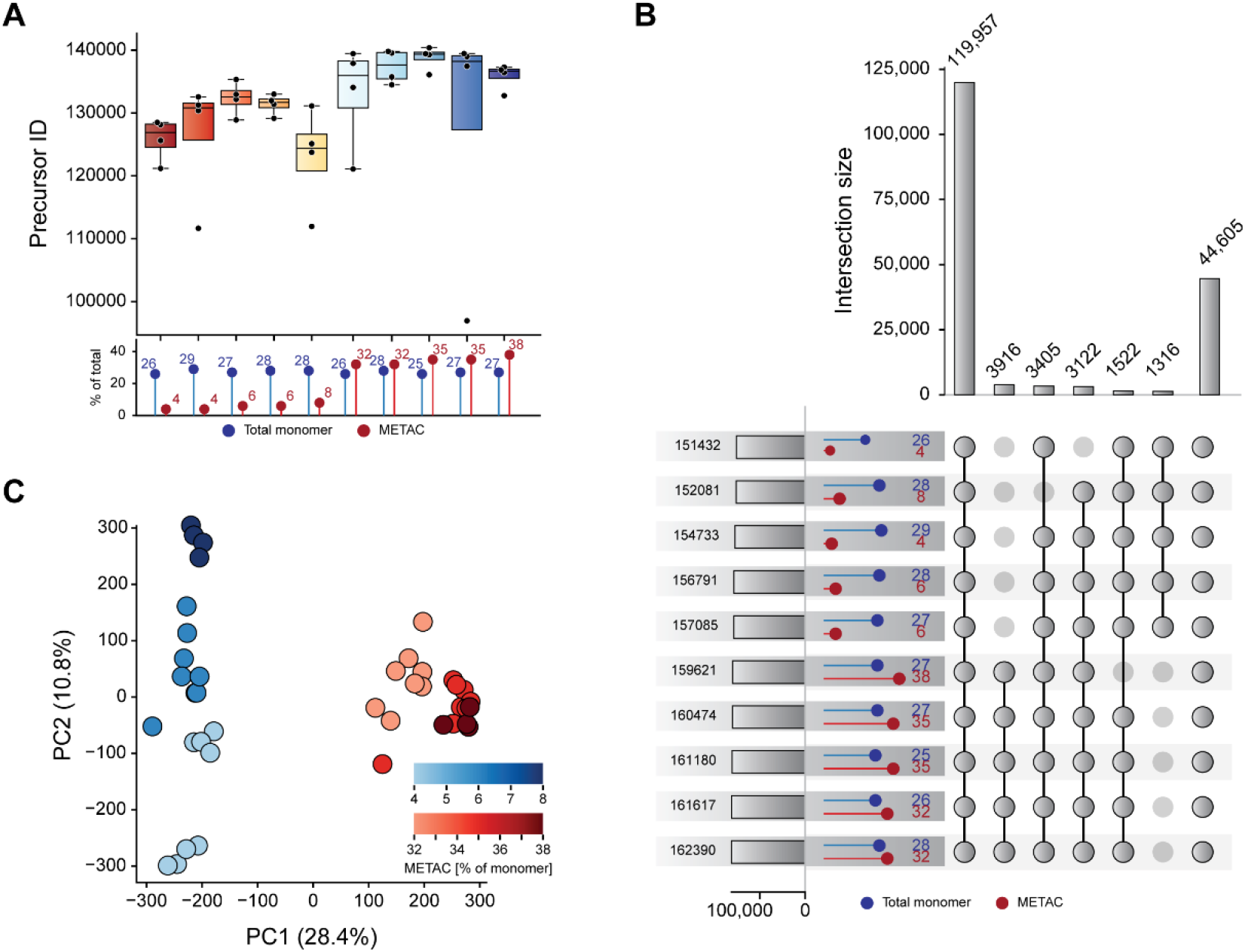
K562 composition screen, precursor level. (A) Precursors per composition (box = IQR, line = median, points = replicates). (B) Precursor overlap across compositions. (C) PCA of log_10_precursor intensities, colored by METAC fraction.

**Supplementary Figure 2.**
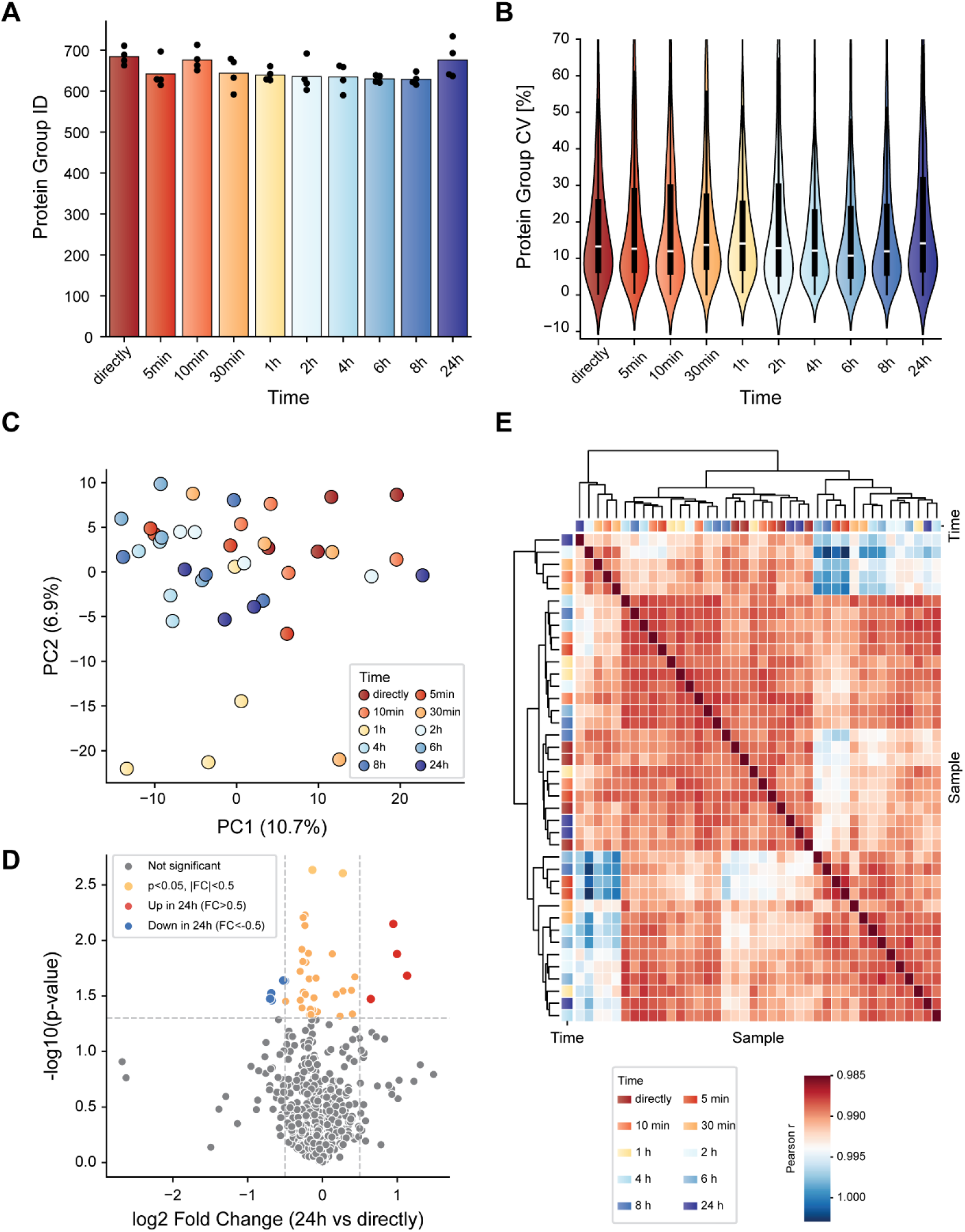
Analysis of phase separation of the precursor mix. A single precursor batch was held in a sealed reservoir at room temperature; six aliquots were withdrawn from a fixed reservoir height at each of ten timepoints after mixing (directly, 5, 10, 30 min, 1, 2, 4, 6, 8, 24 h), polymerized into PP tips on the Bravo platform, and used to process pooled plasma (n = 60 tips total). (A) Protein-group identifications per sample; bars = mean, points = individual replicates. (B) Distribution of within-timepoint protein-group CVs (violin = kernel density, box = IQR, white line = median). (C) Principal-component analysis of log_10_protein-group intensities (complete-case proteins); each point = one tip, colored by time after mixing. (D) Volcano plot of differential abundance between directly-mixed and 24-h-aged precursor; the dashed thresholds mark nominal p < 0.05 and |log_2_FC| > 0.5. (E) Sample–sample Pearson correlation matrix on log_10_protein-group intensities, hierarchically clustered (average linkage on 1 – r).

**Supplementary Figure 3.**
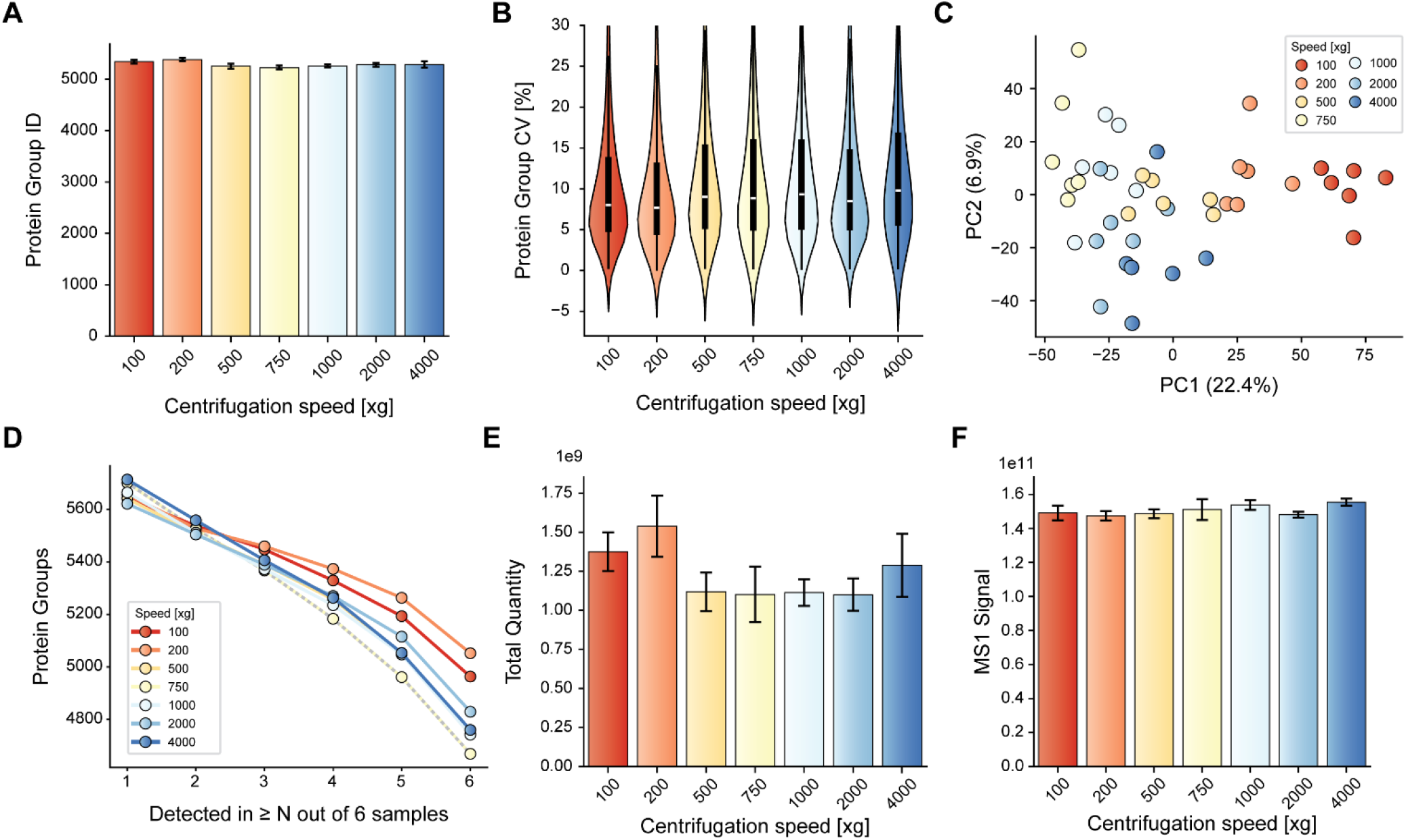
Effect of sample-loading centrifugation speed on SPEC performance at 5 ng K562 lysate input. (A) Protein groups identified per sample (mean ± SD across replicates). (B) Per-protein-group coefficients of variation per condition, computed on raw intensities across the six retained replicates; violins show the full distribution, with the inner box indicating interquartile range and median. (C) Principal component analysis on log10 protein-group intensities (KNN imputation with k = 5, standardized features); each dot represents one sample. (D) Data-completeness curves: number of protein groups detected in ≥ N of 6 replicates per condition; each line represents one centrifugation speed. (E) Total reported peptide quantity per sample (DIA-NN, mean ± SD). (F) Total MS1 signal per sample (DIA-NN, mean ± SD).

**Supplementary Figure 4.**
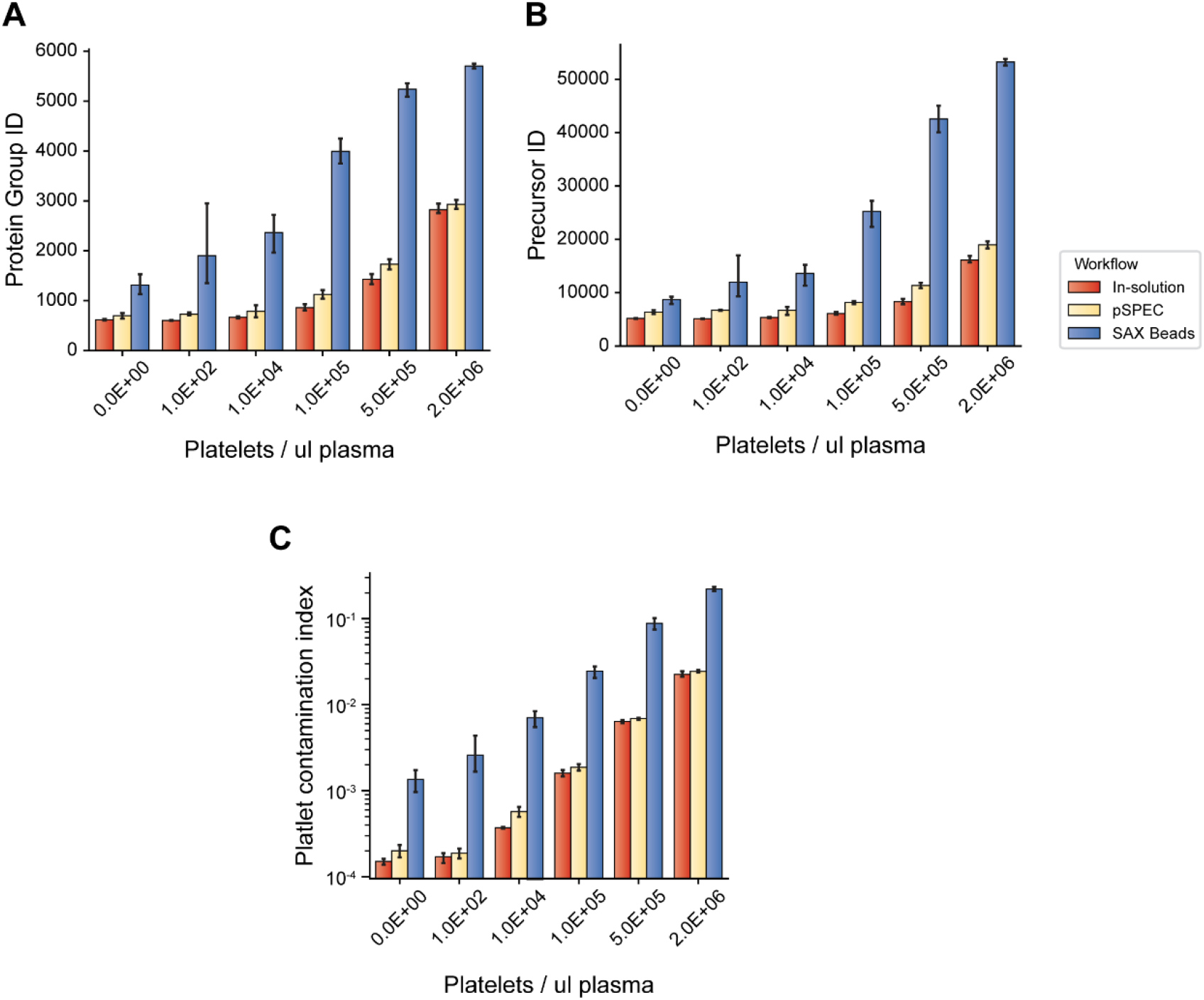
Platelet spike-in contamination controls. (A,B) Protein groups (A) and precursors (B) vs platelet amount. (C) Platelet contamination index vs platelet amount.

**Supplementary Figure 5.**
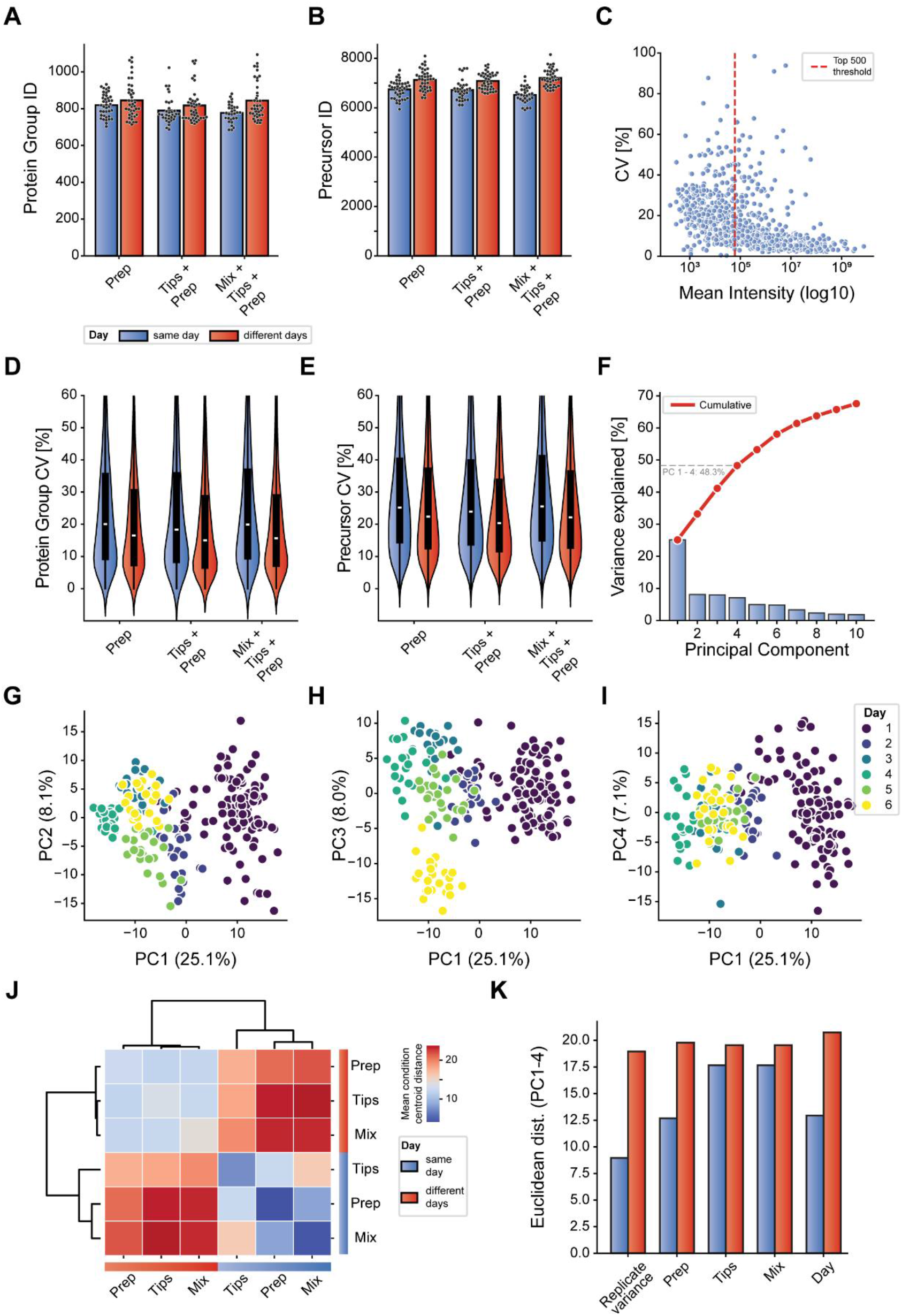
Reproducibility, detailed analyses. (A,B) Protein-group (A) and precursor (B) depth across conditions and days. (C) CV vs protein abundance. (D,E) Tip-to-tip CV, same-day vs different-days, at protein-group (D) and precursor (E) level (violin = density, box = IQR, line = median). (F) PCA scree plot. (G–I) PC1 vs PC2 (G), PC3 (H), PC4 (I), colored by day. (J) Hierarchical clustering of mean condition-centroid distances. (K) Per-factor absolute Euclidean distances.

**Supplementary Figure 6.**
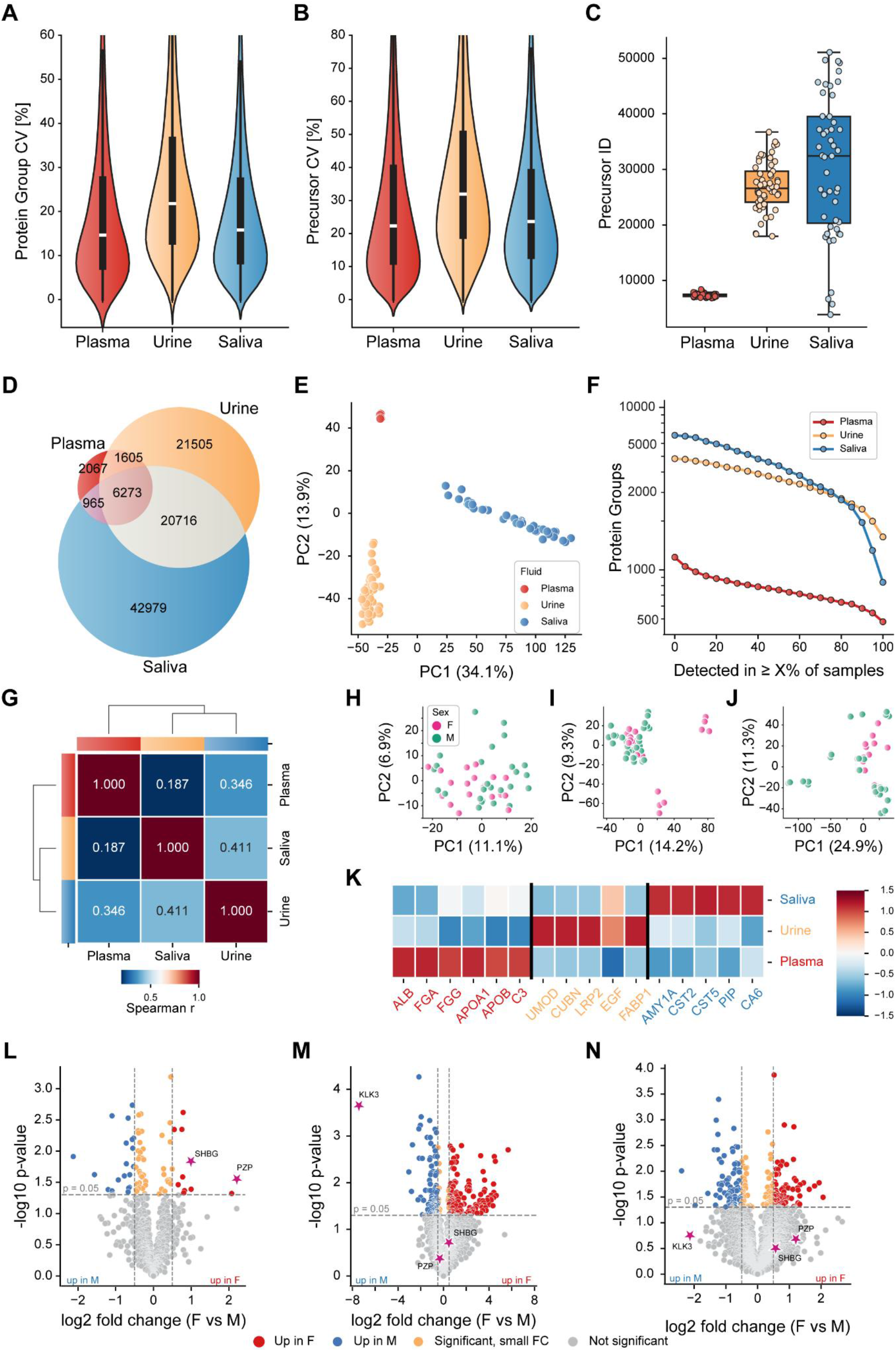
Multi-biofluid cohort: reproducibility, overlap, completeness, structure. (A,B) Per-individual CV across triplicate, by fluid, at protein-group (A) and precursor (B) level (violin = kernel density, box = IQR, white line = median). (C) Venn diagram of precursor overlap across fluids. (D) Protein groups detected in ≥ X% of samples, per fluid (log y). (E–G) Per-fluid PCA of log_10_protein-group intensities (KNN imputation k = 5, z-scored), colored by sex: plasma (E), urine (F), saliva (G). (H– J) Five most abundant proteins per fluid by mean log_10_intensity: plasma (H), urine (I), saliva (J).

